# Cold priming memory reduces plant pathogen susceptibility based on a functional plastid peroxidase system

**DOI:** 10.1101/2020.02.19.956540

**Authors:** Thomas Griebel, Alina Ebert, Hoang Hung Nguyen, Margarete Baier

**Author notes:** **Corresponding Author:** Thomas Griebel, Plant Physiology, Dahlem Centre of Plant Sciences, Freie Universität Berlin, Königin-Luise-Straße 12-16, 14195 Berlin. **Author contributions:** TG, AE, and HHN performed the experiments and analysed the data, TG and MB designed and supervised the research; TG wrote the article and prepared the figures with inputs from MB. All authors read and approved the final version of the article.

## Abstract

Chloroplasts, as recently shown, serve as cold priming hubs in modulating the transcriptional response of *Arabidopsis thaliana* to a second cold stimulus after a stress-free interphase of several days. Here, we studied if such a single 24 h cold stress at 4 °C also alters the susceptibility of Arabidopsis to virulent *Pseudomonas syringae* pv. *tomato* DC3000. Our data show that cold priming did not only increase resistance of Arabidopsis to a subsequent infection immediately, but also after a memory phase of 5 days, during which thylakoid ascorbate peroxidases accumulate. Contrasting to susceptibility, the prior cold exposure did not alter resistance against avirulent and effector-triggered immunity-inducing *Pseudomonas syringae* strains. The effect of cold priming on the plant pathogen susceptibility was independent of the central nucleo-cytoplasmic immune regulator EDS1 (Enhanced Disease Susceptibility 1) and uncoupled from classical immune gene activation. The priming benefits against pathogens required thylakoid and stromal ascorbate peroxidase activity. Combinatorial priming of Arabidopsis pathogen susceptibility by metastable regulation of stromal ascorbate peroxidase activity and post-cold expression of thylakoid ascorbate peroxidase guarantees immediate protection without latency time and prolonged protection by the memory element that regulates future cold responses.

**One-sentence summary:** 24 hour cold exposure reduces plant susceptibility against virulent pathogens dependent on chloroplast ascorbate peroxidases.

## INTRODUCTION

Plants have evolved sophisticated molecular networks that respond differently to simultaneous or sequentially experienced stress events than to single stress situations (Zhang and Sonnewald, 2017; Saijo and Loo, 2019). The concept of priming describes a combination of two sequential and transient stress situations in which the exposure to a prior stress leads to earlier, faster, stronger, and/or more specific responses during the subsequent triggering stress (Crisp et al., 2016; Hilker et al., 2016). Although plants lack a nervous system and an antibody-based adaptive immune system, the plant capacity for a (stress) memory is evident and well described (Conrath, 2011; Crisp et al., 2016; Hilker et al., 2016; Gourbal et al., 2018). The molecular priming memory is formed, while primary stress responses are lost during a stress-free interphase (lag or memory phase), and subsequently modifies the response to a later triggering stimulus (Hilker et al., 2016). The priming memory can result of chromatin modifications, but also be imprinted by preparatory formation or persistence of key signalling metabolites and proteins which are kept in an inactive form during the stress-free memory phases (Conrath, 2011; Crisp et al., 2016; Baier et al., 2019).

The priming and the subsequent stress events can be of same nature (*cis*-priming) or distinct from each other (*trans-*priming) (Hilker et al., 2016). The phenomenon of systemic acquired resistance (SAR) is an intensively studied example of *cis*-priming in plants, in which a pathogen infection leads to an improved immune response in distant, uninfected tissues of the same plant (Conrath, 2011). SAR requires long distance signalling and provides long-lasting protection against a broad range of pathogens (Fu and Dong, 2013; Shah and Zeier, 2013). Several molecular components cooperate in establishing SAR (Conrath, 2011; Fu and Dong, 2013; Shah and Zeier, 2013; Hartmann and Zeier, 2019). These include the co-acting metabolites N-hydroxypipecolic acid, a derivative of the non-proteinogenic amino acid pipecolic acid, and the plant hormone salicylic acid (Hartmann and Zeier, 2019), mitogen-activated protein kinases (MPK) 3 and 6 (Beckers et al., 2009; Wang et al., 2018), the transcriptional co-activator NPR1 (Non-expressor of PR1) (Cao et al., 1997), and the lipase-like protein EDS1 (Enhanced Disease Susceptibility1) (Breitenbach et al., 2014). Pathogen-induced priming leads to broad transcriptional reprogramming in the uninfected plant tissue including chromatin opening and modification (Jaskiewicz et al., 2011; Gruner et al., 2013; Baum et al., 2019).

In contrast to multiple and ternary stress concepts, the dual plant-pathogen interaction based on the innate immune system is broadly and conceptually understood (Jones and Dangl, 2006; Lolle et al., 2020). Plants detect pathogens through recognition of pathogen-associated molecular patterns (PAMPs) by cell surface-exposed pattern recognition receptors (PRRs). PRR activation induces defence responses summarized as PRR- or PAMP-triggered immunity (PTI), which are efficient against a broad range of pathogens (Macho and Zipfel, 2014). Host-adapted and virulent pathogens supress PTI by secreting virulence proteins, so-called effectors, into the host cells with the aim to manipulate cellular physiology and to supress innate immunity (Büttner, 2016). This process strongly affects the susceptibility of the plant and is designated accordingly as effector-triggered susceptibility (ETS) (Jones and Dangl, 2006). A further layer of pathogen defense comprises intracellular nucleotide-binding leucine-rich repeat immune receptors (NLRs) that intercept the presence or activity of pathogen virulence effectors and activate plant responses summarized as effector-triggered immunity (ETI) (Lolle et al., 2020). Two structurally different N-terminal domains, Toll-interleukin1 receptor-like (TIR) and coiled-coiled (CC) form two major groups of plant NLRs: TNLs and CNLs, respectively (Jacob et al., 2013). For instance, the CNL RPM1 (Resistance to *Pseudomonas syringae* pv. *Maculicola* 1) detects the presence of the bacterial effector avrRPM1 by sensing its virulence activity on the RPM1-interating protein 4 (RIN4) (Mackey et al., 2002). Recently, it was shown that ADP-ribosylation on threonine 166 of RIN4 by avrRpm1 precedes and enables its phosphorylation which triggers RPM1-mediated immune signalling (Redditt et al., 2019). An alternative scenario is described by immune activation through the TNL receptor pair RRS1 (Resistance to *Ralstonia Solanacearum* 1) and RPS4 (Resistance to *Pseudomonas syringae* 4) (Griebel et al., 2014). When the bacterial effector AvrRps4 from *Pseudomonas syringae* pv. *pisi* is expressed in the otherwise virulent strain *Pseudomonas syringae* pv. *tomato* DC3000 (hereafter: *Pst* DC3000), ETS is turned into ETI, when RPS4 and RRS1 are present in the infected Arabidopsis accession (Hinsch and Staskawicz, 1996; Wirthmueller et al., 2007; Birker et al., 2009). At the molecular level, avrRPS4 interferes with plant WRKY transcription factors through physical interaction to promote plant susceptibility (Sarris et al., 2015). The TNL pair RPS4/RRS1 traces avrRPS4 interference by using the integrated WRKY domain in RRS1 as a decoy and trap (Le Roux et al., 2015; Sarris et al., 2015). While the bacterial effectors and corresponding NLR pairs inducing ETI are numerous, all TNL receptors share the requirement of the nucleo-cytoplasmic immune regulator EDS1 for all identified signalling responses (Aarts et al., 1998; Wiermer et al., 2005; Wirthmueller et al., 2007; Griebel et al., 2014).

Recently, we have shown that a mild 24 h lasting cold exposure primes cold sensitivity of genes in *Arabidopsis thaliana* for several days (van Buer et al., 2016). Consistent with increased expression of thylakoid-bound ascorbate peroxidase (tAPX) after cold priming, conditional expression of *tAPX* established a cold memory in absence of the priming cold treatment (van Buer et al., 2019). 24 h cold increases specifically expression of stromal ascorbate peroxidase (sAPX) and the gene for the ascorbate regenerating enzyme monodehydroascorbate reductase (MDR; At1g63940 (van Buer et al., 2016). The transcript levels re-attenuate after the stress to control levels within 24 h, while *tAPX* transcript levels increase for 3-4 days after the cold stimulus and tAPX protein accumulates in the thylakoids (van Buer et al., 2016). Our aim is to test for cross signalling effects of cold priming on biotic triggering responses in *Arabidopsis thaliana*. In detail, we investigate if cold priming as the 24 h cold treatment and concomitant specific and subtle modification of the chloroplast antioxidant system are sufficient to improve not only cold-specific signalling but also plant responses against the hemi-biotrophic bacterial pathogen *Pseudomonas syringae*. With transgenic lines or mutants deficient or deprived in the chloroplast ROS detoxification capacities, chloroplasts have been manifold shown to control plant pathogen susceptibility (Sowden et al., 2018). Our data demonstrate that a 24 h long cold priming stimulus reduces susceptibility of *Arabidopsis thaliana* against virulent *Pseudomonas syringae* for at least the 5-7 days. We show that cold priming of pathogen susceptibility is independent of the central immune component EDS1 and key immune gene activation. While the benefits for cold triggering strictly require *tAPX* and are established during the post-cold phase (van Buer et al., 2016; van Buer et al., 2019), cold priming of pathogen susceptibility requires *tAPX* and *sAPX*.

## RESULTS

### 24 h cold exposure reduces plant immune susceptibility for up to five days in an EDS1-independent manner

As shown previously, 4-week-old *Arabidopsis thaliana* Columbia-0 (Col-0) plants memorize a 24 h exposure (including day and night phase) to mild cold (4°C) for up to 7 days (van Buer et al., 2019). Such cold priming leads to modified activation of cold stress-responsive genes during a second cold treatment (van Buer et al., 2016; van Buer et al., 2019). To study, if cold priming and memorization also affects the plant immune response, we triggered cold-pretreated plants (4°C, 24 h) immediately (1 h) or after a memory phase of 5 days by infiltration with the virulent bacterial pathogen *Pseudomonas syringae* pv. *tomato* DC3000 (*Pst* DC3000). Cold-primed Col-0 plants (P, priming) showed significantly reduced *Pst* DC3000 titers (60 %) 3 days after infiltration compared to unprimed control plants (C; control) (Fig. 1). This was not restricted to an early (1h) infiltration time point after cold exposure, but similar strong when a lag phase of 5 days was inserted between cold priming and pathogen triggering (Fig. 1). Consequently, the priming effect of the cold treatment counteracted pathogen growth in the plant not only during the period when cold regulation of gene expression weakens out, but also later when the priming effect on the cold sensitivity is established (van Buer et al., 2019).

**Figure 1.**
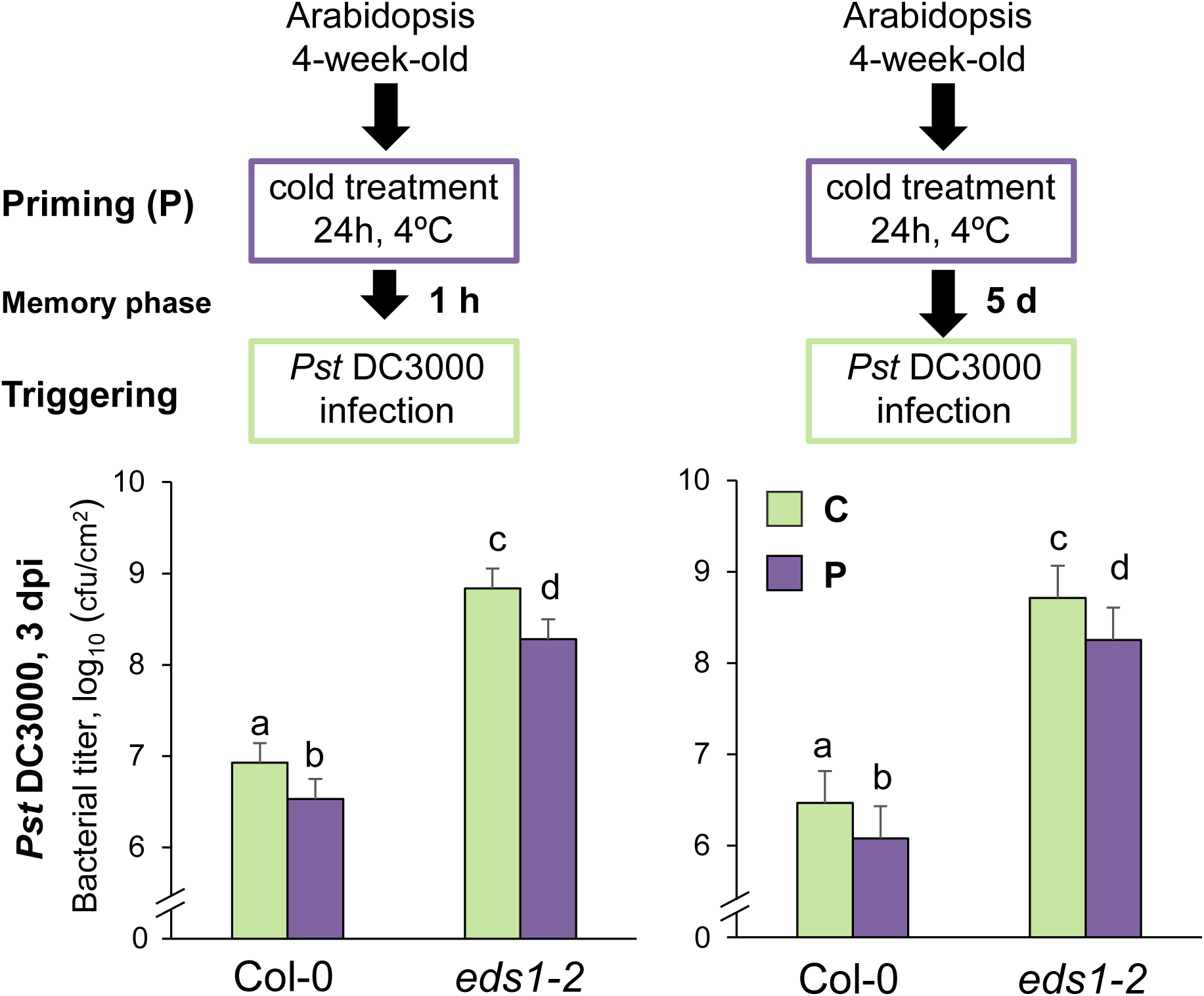
Bacterial growth of *Pst* DC3000 in cold-primed Col-0 and *eds1-2*. 4-week-old *Arabidopsis thaliana* plants were exposed to a 4°C cold treatment of 24 hours (P; Priming). 1 hour (left panel) or 5 days (right panel) after cold exposure, plants were infiltrated with virulent *Pst* DC3000 (OD_600_ = 0.001). Bacterial titers (log_10_-transformed) in cold-primed (P) and control plants (C) of Col-0 and *eds1-2* null mutants were determined 3 days after infiltration (dpi = days post infection). Bars represent means and standard errors calculated from five independent experiments each with 5-6 biological replicates using a mixed linear model. Different letters above the bars denote statistically significant differences (adjusted *p* < 0.05; two-tailed t-tests). cfu/cm^2^, colony forming units per cm^2^ leaf area

To distinguish cold priming regulation from induced basal immune and resistance responses, we included the strongly immune-compromised *eds1-2* null mutant (Bartsch et al., 2006) into our analyses. EDS1 (Enhanced Disease Susceptibility1) is an essential molecular plant immune regulator that contributes to basal resistance and SAR and is required for TNL-mediated ETI (Aarts et al., 1998; Breitenbach et al., 2014; Wirthmueller et al., 2007). The enhanced susceptibility of *eds1-2* against *Pst* DC3000 was significantly reduced when cold-primed plants were inoculated immediately or after 5 days recovery (72 % and 65 % reduction of bacterial titers) (Fig. 1). In these experiments, equal density of starting inoculum was confirmed by measuring bacterial titers 3 hours after plant infiltration ensuring similar starting amounts for bacterial growth rates in control and cold-primed plants (Supplemental Fig. S1). Our data indicate that Arabidopsis benefited with enhanced resistance against virulent *Pst* DC3000 from a 24 h cold priming which was memorized for up to 5 days. Furthermore, cold priming and its memorization weakened plant susceptibility independent or downstream of EDS1-mediated immune signalling.

### Cold response and recovery is functional in immune-impaired *eds1*

Priming and memory concepts require the perception and response of a first (priming) stimulus which initiates the formation of a molecular memory for future stresses (Hilker et al., 2016). We compared the initial cold response and the recovery between Col-0 and *eds1-2* at the transcriptional level of selected genes directly after 24 h exposure to 4°C (0 h), after 72 h, and after 120 h. For this analysis, we harvested plant material only from leaves which we would also have used for bacterial infiltrations. We selected four genes based on recent work on cold *cis-*priming: *COR15A* (*Cold-Regulated Gene15A*; At2g42540), *ZAT10* (*Zinc finger of Arabidopsis Thaliana10*; At2g27730), *BAP1* (*Bon Associated Protein1*; At3g61190), *PAL1* (*Phenyl Ammonia Lyase1*; At2g37040) (van Buer et al., 2016). *COR15A*, which encodes a protein protecting the inner envelope of chloroplasts against freezing damage, is strongly induced during cold exposure and quickly reset (within 24 h) at optimal growth temperatures (Steponkus et al., 2002; Zarka et al., 2003). Same applies to the ROS-induced and pleiotropic stress-responsive genes *ZAT10, BAP1* and *PAL1* (van Buer et al., 2016; van Buer et al., 2019). Cold induction of the selected genes reached a similar level in Col-0 and *eds1-2* at the end of the cold exposure (0 h) and was reset to control rates within 72 h and 120 h (Fig. 2). Basal levels of *ZAT10* and *BAP1* were slightly reduced in *eds1-2* control plants compared to Col-0 (Fig. 2). The conversion of L-phenylalanine to cinnamic acid by PAL1 is a key enzymatic step for a multitude of metabolites such as anthocyanins and flavonoids but also for the synthesis of basal amounts of the plant hormone salicylic acid (SA) (Vlot et al., 2009). However, pathogen-induced SA is mainly metabolized by ICS1 (Iso-Chorismate Synthase 2) and its gene expression is strongly induced after pathogen attack (Hartmann and Zeier, 2019). We could not detect a clear and significant up-regulation of *ICS1* at the end of cold priming exposure and during the subsequent memory phase (Fig. 2). In all our samples, transcripts of SA-responsive immune marker gene *PR1* (*Pathogenesis-Related 1*) remained at low and basal levels and were not detectable. The non-responsiveness of *ICS1* and *PR1* during the post-cold phase distinguishes cold priming-reduced susceptibility from SAR when a first infection leads to immune gene up-regulation (including *ICS1* and *PR1)* in the non-infected systemic tissue (Gruner et al., 2013; Bernsdorff et al., 2016; Hartmann and Zeier, 2019). This is further supported by the functionality of cold-reduced susceptibility in the mutant *eds1-2* (Fig. 1), that is impaired in establishing SAR (Breitenbach et al., 2014). Overall, this analysis suggests that cold signalling during and after exposure to 4°C is perceived and processed similar in Col-0 and in plants lacking EDS1, a central regulator of immunity. While infections directly after cold exposure might benefit from overlapping with post-cold deacclimation of gene expression, infections 5 days after priming require a molecular memory, because cold-induced genes are already reset for at least 2 days.

**Figure 2.**
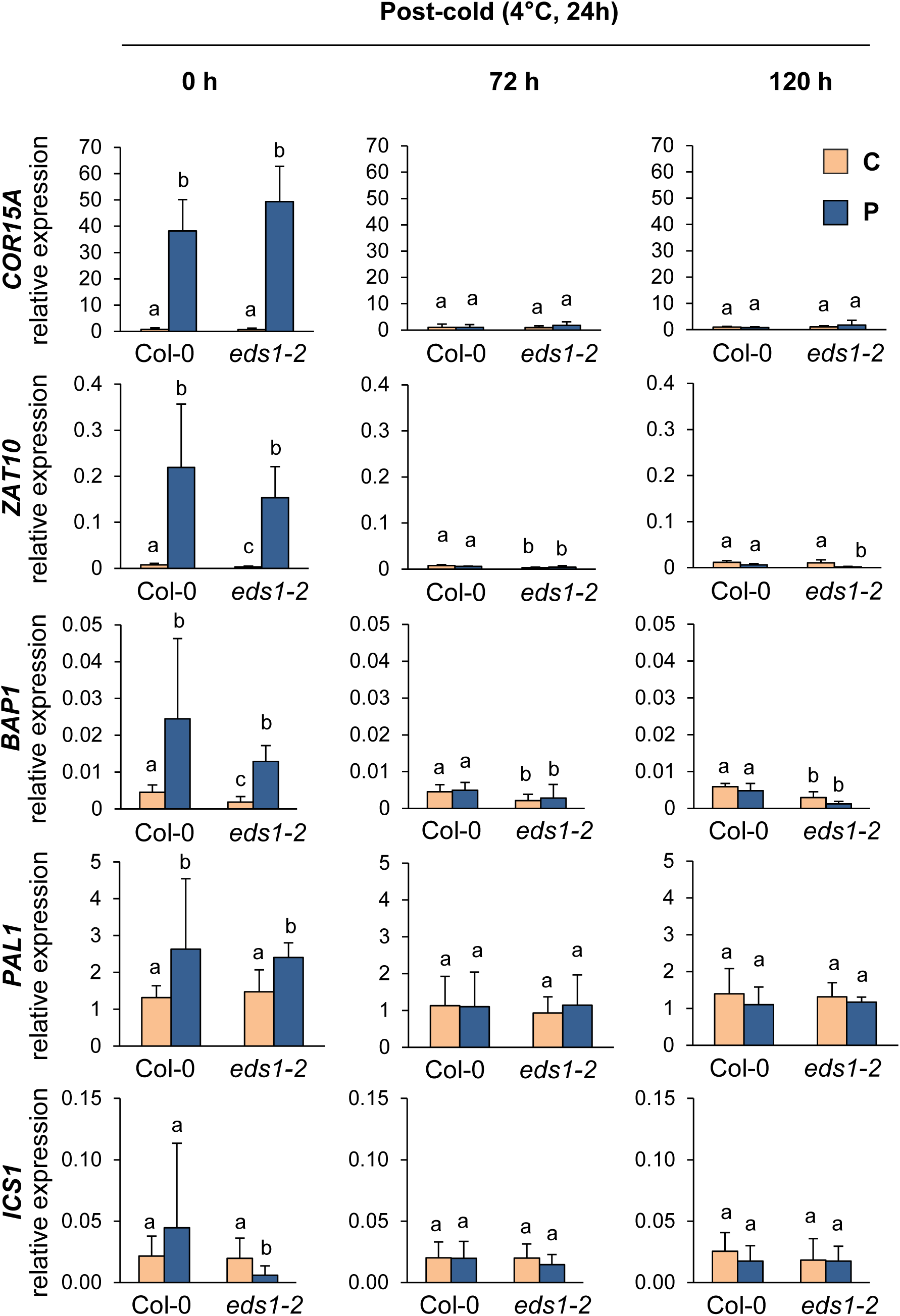
Post-cold expression of stress-responsive genes in Col-0 and *eds1-2*. Transcript levels of *COR15A, ZAT10, BAP1, PAL1*, and *ICS1* in leaves of Col-0 and *eds1-2* null mutants immediately (left panel), 72 hours (middle panel) and 120 hours (right panel) after end of cold priming (P; 4°C, 24 h) were determined by RT-qPCR. Transcript levels in leaves harvested from control plants (C) of same age are shown aside. Bars represent means and standard deviation from three independent experiments as relative expression to the geometric mean of three reference genes (*ACT2, YLS8*, and *RHIP1*). Statistical differences were calculated from log_2_ transformed transcript levels using a mixed linear model followed by two-tailed t tests. Different letters above the bars denote statistically significant differences (adjusted *p* < 0.05).

### Pathogen-induced transcript levels of selected PTI genes and SA signalling are independent from crosstalk with the cold priming memory

The beneficial effect of priming is predicted to result in a faster, earlier, stronger, and/or more sensitive response during the second (triggering) stress (Hilker et al., 2016). We tested transcript levels of four selected genes as indicators for primed activation of classical induced pathogen resistance. Transcriptome dynamics upon infections with virulent *Pst* DC3000 are established rather late between 16 and 24 hours after infection, whereas ETI-inducing pathogens trigger mainly identical transcriptional changes already between 4 and 6 hours (Mine et al., 2018). Hence, we selected 6 and 24 hours after infection for gene expression analysis during pathogen triggering. Expression levels of PTI-inducing genes *NHL10* (*NDR1/HIN1-Like10*, also known as *YLS9*; At2g35980) and MAPK-specific target gene *FRK1* (*FLG22 induced Receptor Kinase 1*; At2g19190) (Boudsocq et al., 2010) were monitored in leaves of cold-primed and control plants after *Pst* DC3000 infection. *NHL10* and *FRK1* were significantly induced 24 hours after infiltration in Col-0, but at a very low basal level in *eds1-2* (Fig. 3). Expression levels in cold-primed and control plants did not differ in time or intensity (Fig. 3). We tested for cold priming-responsive expression profiles of SA-biosynthetic *ICS1* and the SA-responsive *PR1. ICS1* and *PR1* were significantly induced at 24 h, but not 6 h after the pathogen treatment in Col-0 (Fig. 3). The induction level or the dynamic of the responses did not differ between cold-primed and control samples (Fig. 3) indicating that pathogen-triggered SA-production and signalling is not cold-primed and thereby not causative for the cold-reduced susceptibility. This is further supported by the requirement of functional EDS1 for a robust activation of SA-related immune pathways upon infection with virulent pathogens (Rietz et al., 2011; Cui et al., 2017). Cold priming does not affect *ICS1* and *PR1* levels downstream or independent of EDS1 as induced transcripts in Col-0 are absent or low in cold-pretreated and inoculated *eds1-2* (Fig. 3).

**Figure 3.**
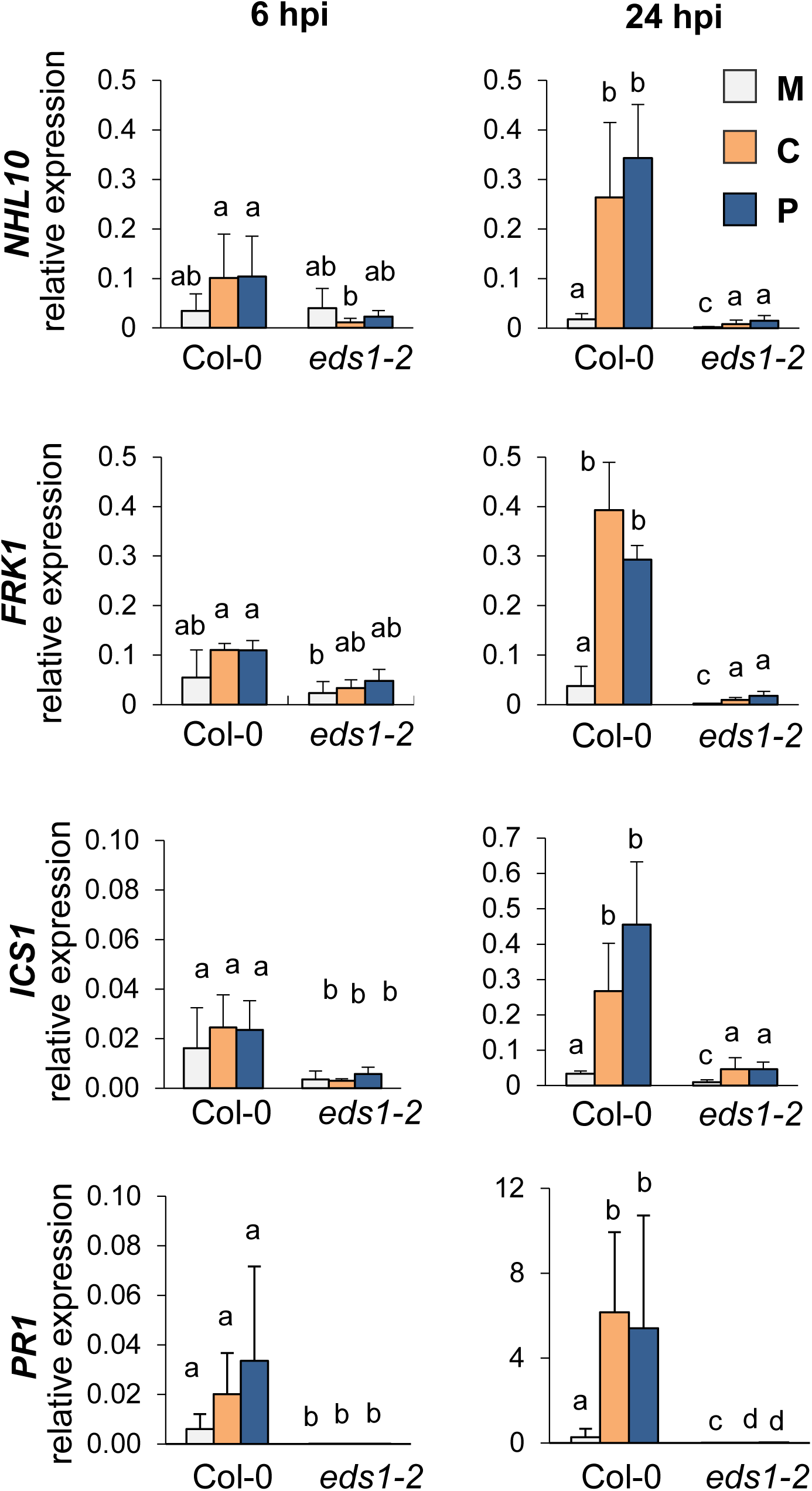
Transcript levels of selected pathogen-responsive genes during *Pst* DC3000 infection upon cold priming. Transcript levels of PTI marker genes *NHL10* and *FRK1*, SA biosynthetic *ICS1* and SA-responsive *PR1* in leaves of Col-0 and *eds1-2* null mutants after infiltration with *Pst* DC3000 (OD_600_ = 0.005). Infiltrations were done 5 days after end of cold treatment using cold-primed plants (P; 4°C, 24 h) or control plants (C). Transcript levels in mock (M)-infiltrated leaves from control plants of same age are shown aside. Transcript levels were determined 6 (left panel) or 24 (right panel) hours post infection (hpi). Bars represent means and standard deviations from three independent experiments relative to the geometric mean of three reference genes (*ACT2, YLS8*, and *RHIP1*). Statistical differences were calculated from log_2_ transformed expression levels using a mixed linear model followed by two-tailed Student’s t tests. Different letters above the bars denote statistically significant differences (adjusted *p* < 0.05).

### Post-cold and *Pst* DC3000-triggered regulation of *tAPX* and *sAPX* transcripts

Recently, the thylakoid-bound ascorbate peroxidase (tAPX) was described for its role in establishing the memory that controls cold regulation of gene expression after cold priming for several days (van Buer et al., 2016; van Buer et al., 2019). Hereby, post-cold accumulation of *tAPX* transcripts and proteins was essential for the memory function (van Buer et al., 2016; van Buer et al., 2019). In addition to tAPX, Arabidopsis expresses stroma-localized ascorbate peroxidases (sAPX) (Ishikawa and Shigeoka, 2008), which evolved from the same ancesteral gene as *tAPX* and still has a highly similar catalytic subunit (Pitsch et al., 2010). We compared regulation of *tAPX* and *sAPX* after cold exposure and later on after *Pst* DC3000 infection in leaves of cold-primed and control plants of Col-0 and *eds1-2* (Fig. 4). Our data confirm previously described post-cold induction of *tAPX* in Col-0 (van Buer et al., 2016) and show that this memory phase lasting process was also functional and significant in *eds1-2* (Fig. 4a). *sAPX* was up-regulated during a 24 h cold phase of 4 °C (van Buer et al., 2016) (Fig. 4a) and quickly adjusted to pre-cold levels after the cold treatment. Similar to cold, infiltration of leaves with *Pst* DC3000 reduced *tAPX* levels in Col-0 and *eds1-2* between 3 and 24 hours after infection in naïve and cold-pretreated plants (Fig. 4b). Pathogen-induced *sAPX* activation in Col-0 was completely absent in *eds1-2* and independent of a prior cold priming (Fig. 4b). Thereby, our data reveal a regulatory similarity between cold- and pathogen-responsive cellular plant stress management: up-regulation of *sAPX* (EDS1-dependent under pathogen stress) and repressive regulation of *tAPX* (EDS1-independent). Decisive post-cold induction of *tAPX* during memory phase (van Buer et al., 2016; van Buer et al., 2019) in Col-0 and *eds1-2* further supports EDS1-independent cold priming memory formation.

**Figure 4.**
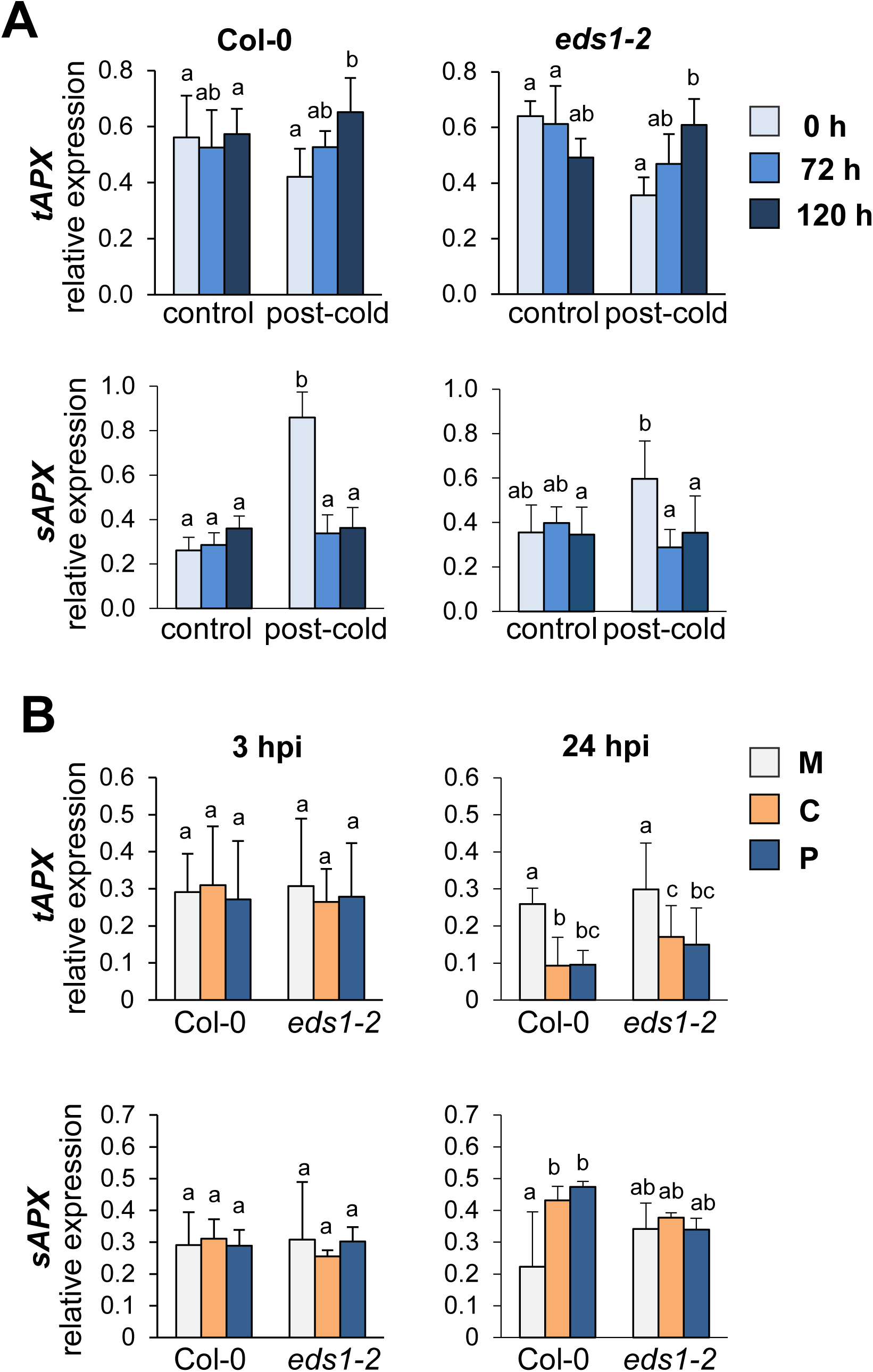
Transcript regulation of plastid ascorbate peroxidases encoding genes *tAPx* and *sAPx* during post-cold phase and after *Pst DC3000* infection. (A) Post-cold (4°C, 24 h) transcript levels of *tAPx* and *sAPx* in leaves of Col-0 (left panel) and *eds1-2* null mutants (right panel) immediately (0 h), 72 hours, and 120 hours after end of cold treatment. Transcript levels in leaves harvested from control plants of same age are shown aside. (B) Transcript levels of *tAPx* and sAPx in leaves of Col-0 and *eds1-2* null mutants 3 hours (left panel) and 24 hours (right panel) post infection (hpi) with *Pst* DC3000 (OD_600_ = 0.005) of cold-primed (P) or control plants (C). Plants were infected 5 days after end of cold treatment. Transcript levels in leaves harvested from mock-infiltrated control plants (M) are shown aside. (A,B) Bars represent means and standard deviation from three independent experiments as relative expression to the geometric mean of three reference genes (*ACT2, YLS8*, and *RHIP1*). Statistical differences were calculated from log_2_ transformed transcript levels using a mixed linear model followed by two-tailed Student’s t tests. Different letters above the bars denote statistically significant differences (adjusted *p* < 0.05).

### Cold priming-reduced pathogen susceptibility is impaired in *tAPX-* and *sAPX*-KO lines

To test if *tAPX* is required not only for cold priming of ROS-responsive genes during cold triggering (van Buer et al., 2019), but also for beneficial responses during *Pst* DC3000 infections, we included *tapx* and sapx knockout lines (Kangasjarvi et al., 2008) into our analysis. Cold-primed (4°C, 24 h) and control plants were infiltrated with *Pst* DC3000 5 days after the priming stimulus (Fig. 5). The bacterial titers measured in control plants revealed that *tAPX* did not *per se* contribute to the degree of plant pathogen susceptibility and basal resistance (Fig. 5), whereas *sapx* KO-lines were slightly but not significantly more resistant against *Pst* DC3000 (Fig. 5). Cold-pre-treated *tapx-*KO and *sapx*-KO lines did not show the priming effect observed in Col-0 plants (Fig. 5). While cold triggering responses specifically required *tAPX*, but not *sAPX* (van Buer et al., 2016; van Buer et al., 2019), cold-primed pathogen susceptibility benefited from functionality of both plastid ascorbate peroxidase variants. Consequently, the memory reducing pathogen susceptibility can be postulated to be more generally H_2_O_2_-controlled than priming of the cold responsiveness.

**Figure 5.**
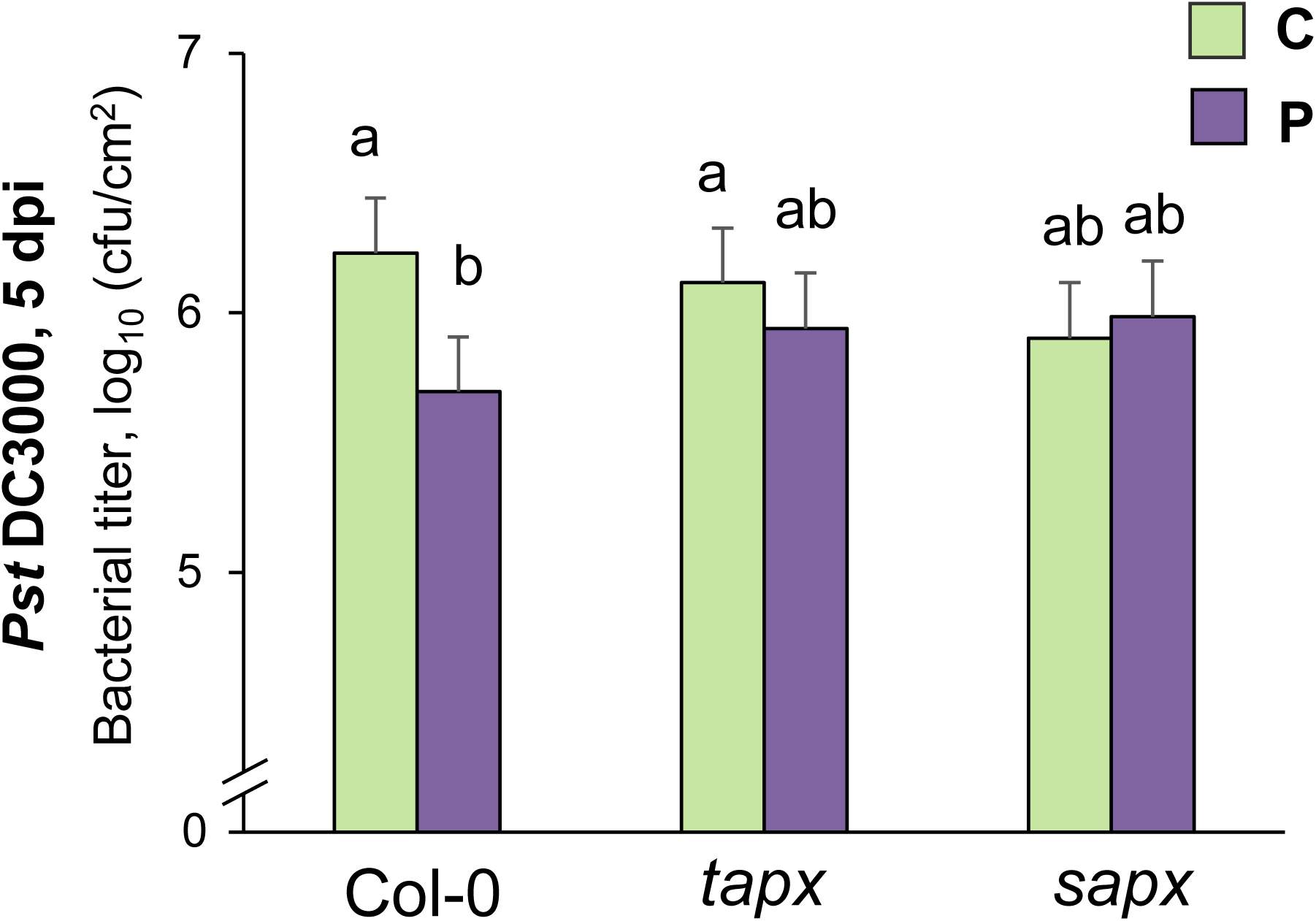
Bacterial growth of *Pst* DC3000 in cold-primed Col-0, *tAPx* and *sAPx* KO lines. 5 days after cold exposure (4°C, 24h) leaves of cold-treated (P) and control (C) plants were infiltrated with *Pst* DC3000 (OD_600_ = 0.001). Bacterial titers (log_10_-transformed) were measured 3 days after infiltration (dpi = days post infection). Bars represent means and standard errors calculated from five independent experiments each with 5-6 biological replicates using a mixed linear model. Different letters above the bars denote statistically significant differences (adjusted *p* < 0.05; two-tailed t tests). cfu/cm^2^, colony forming units per cm^2^ leaf area

### Growth of ETI-activating *P. syringae* is not modulated by prior cold exposure, but virulence activity of bacterial avrRPS4 counteracts cold priming

Plant-pathogenic bacteria, including the virulent *Pst* DC3000 deliver a repertoire of bacterial type III effectors into plant cells which suppresses basal plant defence responses and PTI (Lindeberg et al., 2012; Büttner, 2016). To test if cold priming effects can also be observed on bacterial growth, when bacterial effectors are detected *in planta* and a strong ETI is activated, we used two *Pst* DC3000 strains delivering either the effector avrRPM1 or avrRPS4 (Grant et al., 1995; Hinsch and Staskawicz, 1996). In Arabidopsis Col-0 accession, molecular virulence activity of avrRPM1 is detected by the intracellular CNL receptor complex RPM1 (Resistance to *Pseudomonas syringae* pv. *Maculicola* 1) in an EDS1-independent manner (Aarts et al., 1998; Mackey et al., 2002). Bacterial growth of *Pst avrRPM1* was limited in naïve control plants of Col-0 and *eds1-2* to a similar level and we could not detect differences between cold-primed and control plants after immediate (1 h) or 5-days-later performed inoculations. This indicates that cold priming and memory affects susceptibility rather than resistance initiated through mechanisms of ETI.

The bacterial effector avrRPS4 is recognised *in planta* by the TNL receptor pair RPS4 and RRS1 (Wirthmueller et al., 2007; Saucet et al., 2015). Activation of TNLs converges on EDS1 for all downstream signalling outputs (Griebel et al., 2014). While *Pst avrRPS4* triggers ETI in Col-0, *eds1* remains susceptible (Aarts et al., 1998; Wirthmueller et al., 2007). This was consistent with our measurements of bacterial titers in naïve control plants of Col-0 and *eds1-2* (Fig. 6). Like in *Pst avrRPM1*-infiltrated plants, growth of the ETI-inducing *Pst avrRPS4* in Col-0 was not altered by an earlier cold treatment (Fig. 6). Surprisingly, the susceptibility of *eds1-2* challenged with *Pst avrRPS4* was not reduced after cold treatment (Fig. 6) as it was after *Pst* DC3000 infection (Fig. 1). This indicates that the virulence activity of avrRPS4 antagonizes mechanisms enabling reduced susceptibility after cold priming memory formation.

**Figure 6.**
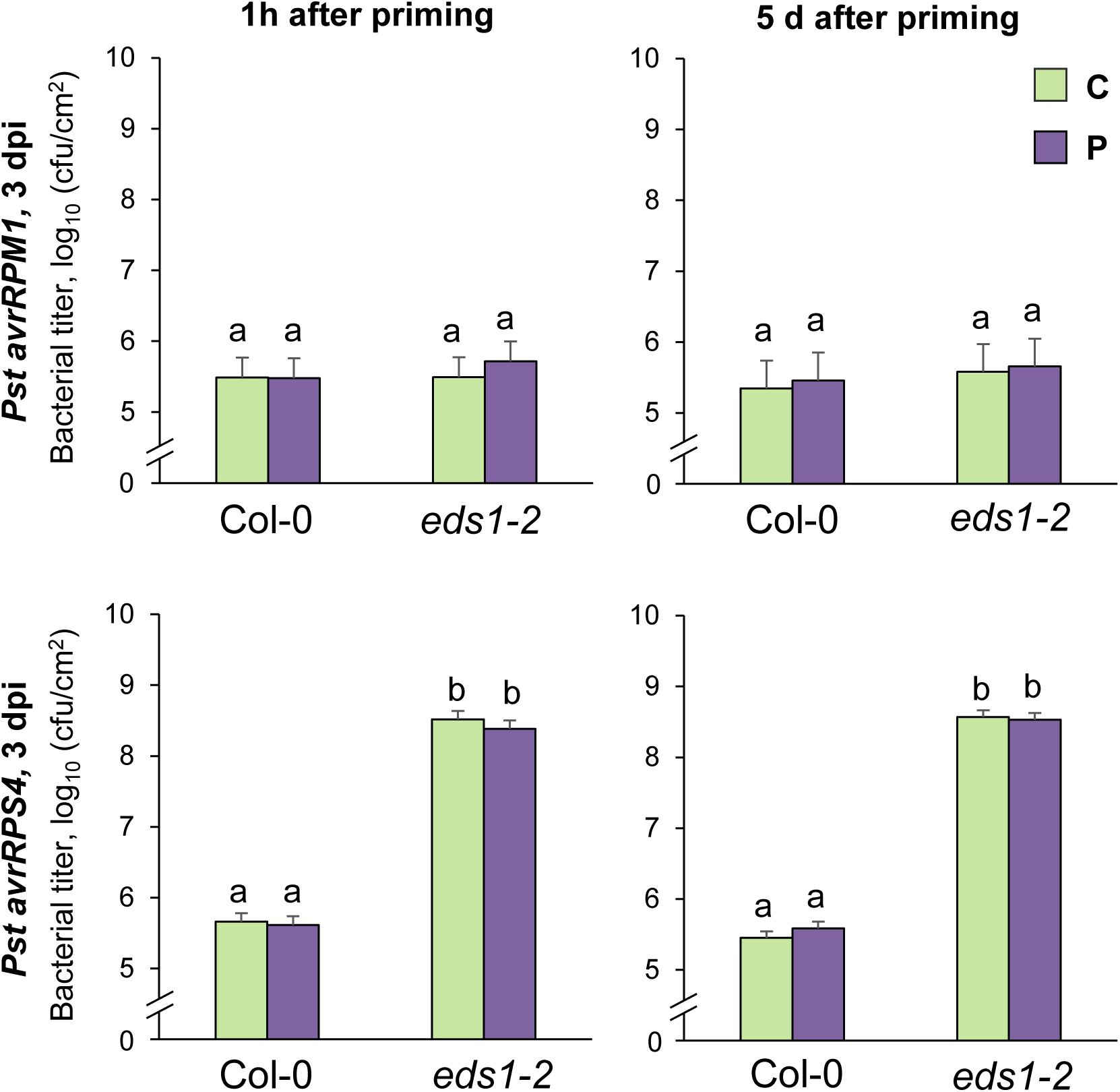
Bacterial growth of *Pst avrRPM1* and *Pst avrRPS4* in cold-primed Col-0 and *eds1-2*. Leaves of cold-primed (4°C, 24h, P) and control (C) plants were infiltrated (OD_600_ = 0.001) with *Pst avrRPM1* (upper panel) or *Pst avrRPS4* (lower panel) after 1 hour (left panel) or 5 days after cold exposure (right panel). Bacterial titers were measured 3 days after infiltration (dpi = days post infection). Bars represent means of log_10_-transformed titers and standard errors calculated from four independent experiments each with 5-6 biological replicates using a mixed linear model. Different letters above the bars denote statistically significant differences (adjusted *p* < 0.05; two-tailed t tests). cfu/cm^2^, colony forming units per cm^2^ leaf area

## DISCUSSION

When plants are exposed to simultaneously or sequentially occurring combined abiotic and biotic stress situations, responses often differ compared to single and individual stresses (Zhang and Sonnewald, 2017). The outcome of different combined stresses can result in a trade-off situation or enable cross-tolerance (Saijo and Loo, 2019). Cross-tolerance upon two sequentially applied stresses including a stress-free lag phase, which enables recovery and requires memorization of the first stressor, is a characteristic feature of the priming phenomenon (Hilker et al., 2016). Here, we showed that a single 24 h cold exposure primed the susceptibility of *Arabidopsis thaliana* Col-0 against the virulent plant pathogen *Pseudomonas syringae* pv. *tomato* DC3000 for up to five days (Fig. 1, Fig. 5). This cold priming-reduced pathogen susceptibility was independent from plant immunity pathways controlled by EDS1, but it required the chloroplast-located ascorbate peroxidases sAPX and tAPX (Fig. 7).

**Figure 7.**
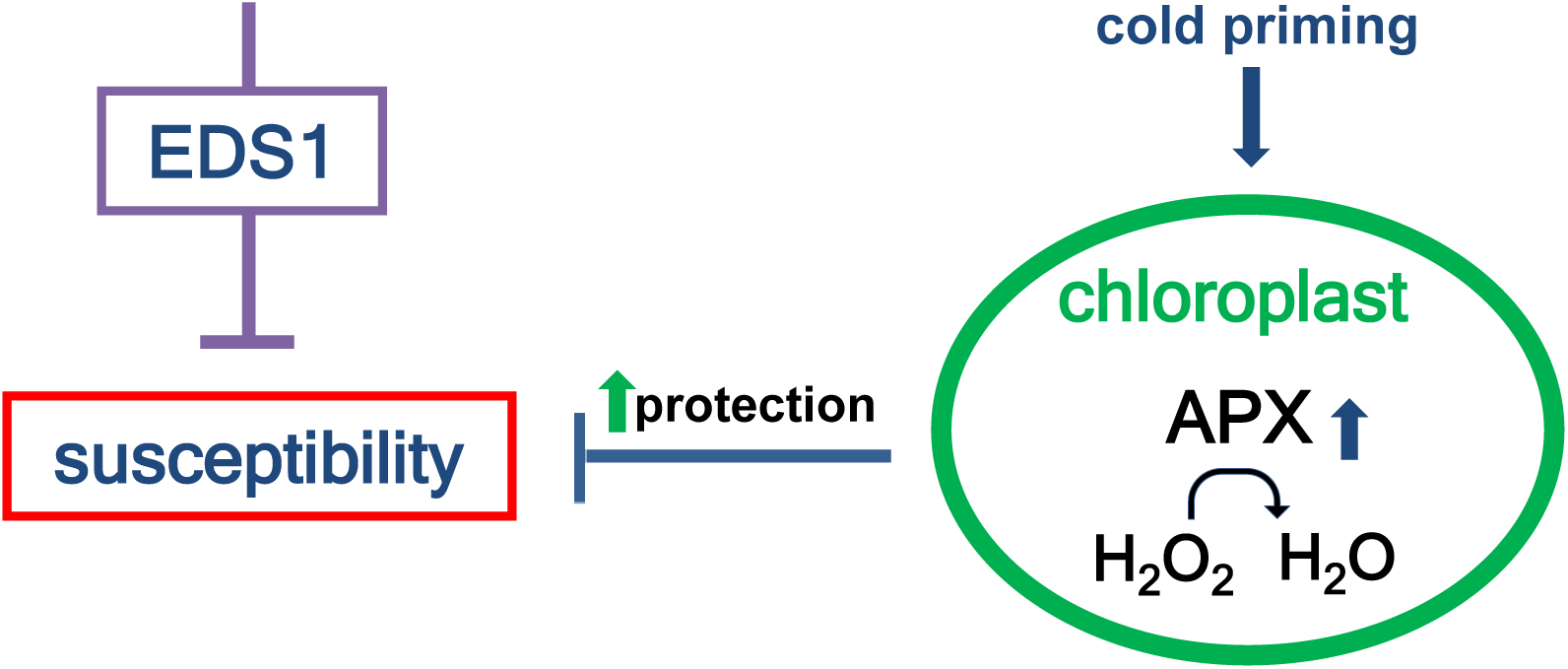
Proposed model for the role of plastid ascorbate peroxidases in reduced plant pathogen susceptibility by cold priming memory. A 24 h lasting exposure to cold (4°C) reduces plant susceptibility in *Arabidopsis thaliana* Col-0 against virulent *Pst* DC3000. This depends on a cold memory that signals through chloroplast stromal and thylakoid ascorbate peroxidases (APX). The chloroplast cold priming effect is independent and additive to the function of the plant immune regulator EDS1 in controlling plant susceptibility.

### Cold priming memory and pathogen susceptibility

The immune system of *Arabidopsis thaliana* benefits from the experience of cold temperatures. A single 24 h cold (4°C) exposure lead to immediate decrease in pathogen susceptibility (shown by infection 1 h after cold), but was also robust for 5-8 days (shown by infection 5 d after cold; Fig 1), although the initial transcriptional cold response (e.g. *COR15A, ZAT10, BAP1, PAL1*) was quickly re-set to pre-cold levels (Fig. 2). As shown recently by Wu et al. (2019), a shorter 10-h cold treatment applied during the night also decreases pathogen susceptibility in *Arabidopsis thaliana*. However, the shorter night stimulus is only transiently memorized for up to 12 h (Wu et al., 2019). Repetitive application of seven 1 h cold periods (1 treatment per day) equally results in reduced growth of *Pst* DC3000 and entrained protection for 7 days (Singh et al., 2014). Our study showed that already a single 24 h lasting cold treatment primed and established a molecular stress memory lasting for 5-8 days. Whereas repetitive cold treatments resulted in enhanced activation of PTI-responsive genes *FRK1* and *NHL10* upon triggering with type-III-secretion-deficient *Pst hrcC* strain (Singh et al., 2014), the single 24 h cold priming stimulus did not reveal a priming pattern for *FRK1* and *NHL10* transcripts. Activation of *FRK1* and *NHL10* without priming signature excludes a similar cross stress memory formation as with the repetitive cold stimuli and shows a PTI-independent memory (Fig. 3).

### Independency of the immune regulator EDS1

Cold priming memory formation is independent from the central plant immune regulator EDS1: (i) bacterial growth was reduced in cold-primed *eds1* null mutants (Fig. 1), (ii) initial cold sensing in *eds1* was wild type-like leading e.g. to *COR15A, ZAT10, BAP1* and *PAL1* activation (Fig. 2), EDS1-dependent transcript activation of selected immune-related genes (*NHL10, FRK1, ICS1, PR1*) did not show a cold priming signature or an activation downstream of EDS1 (Fig. 3). Several EDS1-dependent responses have been well-described for their effects in reducing plant susceptibility to virulent pathogens: EDS1, together with its homolog and heteromeric complex partner PAD4, boosts SA biosynthesis (Cui et al., 2017) and promotes tocopherol production in an SA-independent manner upon *Pst* DC3000 infection (Stahl et al., 2019). Furthermore, EDS1 is required for the plant immune *cis*-priming SAR by contributing to signal generation in primary infected leaves and signal perception in the systemic uninfected tissues (Breitenbach et al., 2014). Based on the functionality of cold priming-reduced susceptibility in *eds1-2*, we conclude that the molecular mechanisms of cold priming memory formation are independent from EDS1-controlled immune activation and established irrespective of SA production during *Pst* DC3000 triggering or SAR signalling (Fig. 7). In addition, the indispensable requirement for EDS1 in TNL-mediated immunity (Wiermer et al., 2005; Griebel et al., 2014) excludes the possibility that (post-)cold activation of TNL immune receptor signalling is causative for the reduced susceptibility in cold-primed plants. Several articles describe a correlation of induced SA under low temperature conditions. As during pathogen attack, cold-induced SA derives from the plastid isochorismate pathway, but SA levels are not increasing before 7 days at 4°C (Kim et al., 2013; Kim et al., 2017). A 24h lasting cold exposure was not sufficient to significantly increase *ICS1* transcript levels (Fig. 2) or enhance SA levels but requires a constant cold exposure of at least one week (Kim et al., 2013; Wu et al., 2019). In contrast to the independency of EDS1 during mild cold exposure (4°C, 24h) and cold memory formation, induction and activation of the EDS1 complex at temperatures below 4°C negatively affect freezing tolerance in an SA-dependent manner (Chen et al., 2015).

### The role of chloroplasts

Chloroplasts can be considered as the cellular origin of cold sensing and priming: cold exposure supports chloroplast ROS production by an imbalance between Calvin-Benson cycle and photosynthetic electron transport (Ensminger et al., 2006) and leads to reduced activation of chloroplast-to-nucleus ROS signalling sensitive genes during a second cold phase (van Buer et al., 2016; van Buer et al., 2019). tAPX, the main regulator of cold priming memory in chloroplasts, is transcriptionally activated during post-cold phase (van Buer et al., 2016; van Buer et al., 2019) in an EDS1-independent manner (Fig. 4). Gene expression regulation upon cold triggering specifically benefits from thylakoid-bound APX activation, but not from the stromal APX (van Buer et al., 2019). However, robust cold priming against virulent pathogens required both forms of plastid ascorbate peroxidases (Fig. 5). Bacterial growth rates in *tapx-KO* were wild type-like and not affected, except for the missing cold memory response (Fig. 5). Similar to cold exposure, infections with virulent pathogens, resulted in reduced *tAPX* transcript levels (Fig. 4). Whereas cold priming of cold triggering is solely regulated by tAPX (van Buer et al., 2016; van Buer et al., 2019), pathogen triggering of cold-primed plants also required functional sAPX for memory effects (Fig. 4). Such regulation indicates a general function of plastid APX in controlling stress-responsive redox state and ROS generation.

Several bacterial effector proteins aim to alter chloroplast functions to increase susceptibility of the plant (Serrano et al., 2016). A chloroplast-located virulence activity is suggested for the bacterial effector avrRPS4 (Li et al., 2014). However, avrRPS4 targets also nuclear WRKY transcription factors through direct interaction as part of its virulence function (Sarris et al., 2015). In absence of functional EDS1, the bacterial effector avrRPS4 is not detected by the plant immune system but clearly contributes to the virulence of the pathogen (Wirthmueller et al., 2007). In our study, we could not observe a cold priming-dependent reduction of *Pst avrRPS4* growth in *eds1* (Fig. 6). Hence, the virulence activity of avrRPS4 either counteracts the cold priming effect or is not dampened through cold-primed cellular changes.

Pathogenic *Pseudomonas syringae* effectors actively suppress plant photosynthesis by lower activity of photosystem II resulting in decreased thylakoid electron transport (de Torres Zabala et al., 2015). This prevents PTI-induced plastid ROS production and ROS-initiated defence responses (de Torres Zabala et al., 2015; Serrano et al., 2016). In addition to counteracting H_2_O_2_ accumulation, cold priming effects on APX availability can render photosystem function and electron transport to be less vulnerable for the manipulation of bacterial effectors. Plastid APX control, for instance, electron flux into the cyclic electron transport and plastoquinone reduction (Asada, 1999; Maruta et al., 2016), which affects plastid function and plastid-to-nucleus signalling. Consistently, pathogen growth is delayed in plants with constitutive overexpression of *tAPX* and correlates with reduced disease symptoms and cell death (Yao and Greenberg, 2006).

### Benefit of connectivity between cold priming and pathogen susceptibility

Concerning the ecological relevance of cold priming-reduced pathogen susceptibility, a central question remains: Is the connectivity of cold priming and pathogen susceptibility part of a specific-adaptive trait for *Arabidopsis thaliana* or result of a general and pleiotropic stress response? Singh et al. (2014) show that in case of repetitive abiotic stress treatments the nature of the environmental stress (e.g. heat, cold, or salt) is not decisive for the primed pathogen resistance indicating that cold priming reduced pathogen susceptibility is generally (co-)activated by fluctuating environmental stress conditions. Reduced temperatures might still indicate a higher risk for pathogen attack due to its benefits for the lifestyle of some relevant Arabidopsis pathogens (Ruhe et al., 2016; Herlihy et al., 2019). Specifically, the successful colonization of *Arabidopsis thaliana* by its natural oomycete pathogen *Hyaloperonospora arabidopsidis* requires decreased cool temperatures (∼16°C) (Herlihy et al., 2019) and for experimental infections with the pathogenic oomycete *Albugo laibachii, Arabidopsis thaliana* plants are kept in the cold for one night phase to promote germination of the spores (Ruhe et al., 2016). Our study demonstrated that a single short (24 h) 4°C phase is sufficient to decrease plant susceptibility against pathogens for several days. This demonstrates that plants evolved strategies to counteract cold- or chilling-linked improved pathogenesis of certain microbes. This several day active mechanism might be of high importance during seasonal times when plants and pathogens face short cold periods with unpredictable duration and frequency. In contrast to cold sensitivity, which requires 2-3 days after the initial cold priming for activation, the combination of immediate and long-lasting cold priming effects on plant immunity ensure efficiency and robustness.

## MATERIALS AND METHODS

### Plant material and cultivation

*Arabidopsis thaliana* var. Col-0 plants, *eds1-2* null mutants (Bartsch et al., 2006), and T-DNA knockout lines *tapx* and *sapx* (Kangasjarvi et al., 2008) were used in this study. All mutant lines are in Col-0 background. Plants were cultivated in round pots (6 cm diameter) containing a soil mixture (14:14:5) of “Topferde” (Einheitserde, Sinntal-Altengronau, Germany), “Pikiererde” (Einheitserde, Sinntal-Altengronau), and, “Perligran Classic” (Knauf, Germany) supplemented with 0.5 g l^-1^ dolomite lime (Deutsche Raiffeisen-Warenzentrale, Germany) and grown in a controlled environmental chamber with a day/night temperature of 20 ± 2 °C and 18 ± 2 C, 10 h light (100-110 µmol photons*m^-2^*s^-1^) and 14 h dark cycle, and a constant relative humidity of 60 %± 5 % after a stratification at 4°C for two days.

### Cold treatments

Cold treatments were performed as previously described (van Buer et al., 2016; van Buer et al., 2019). 4-week-old plants were exposed to cold 2.5 hours after onset of light by transferring them to a growth chamber with constant 4 ± 2°C but otherwise identical aeration, illumination and air humidity as in the 20°C chamber. After a continuous cold exposure for 24 h (comprising a full day and night phase), the plants were placed back to the 20 °C chamber, labelled and randomized with the non-cold-treated control plants.

### Cultivation and inoculation of bacteria

*Pseudomonas syringae* pv. *tomato* DC3000 (*Pst* DC3000), *Pst* DC3000 carrying either the avirulence gene *avrRpm1* (*Pst avrRPM1*) or *avrRPS4* (*Pst avrRPS4*) were grown for 24 h at 28°C on NYGA solid medium containing the appropriate antibiotics. Bacterial cultures were suspended in 10 mM MgCl_2_ and diluted to optical densities of 0.001 at 600°nm (OD_600_) for bacterial growth assays or 0.005 for gene expression analyses. The bacterial suspensions were infiltrated from the abaxial side into the leaves with a needleless syringe. For transcript analyses, control plants were mock (M)-treated with 10 mM MgCl_2_. Bacterial inoculations were performed 3.5 h ± 0.5 h after onset of either 1h or 5 days after the end of cold treatment as indicated. The three youngest, but fully-grown leaves of each plant were selected for infiltration.

### Bacterial growth assays

*In planta* bacterial titers were determined at the indicated time point after infiltration by combining three leaf discs for one sample and shaking in 10 mM MgCl_2_ with 0,01 % (v/v) Silwet L-77 at 28°C for 1 h. From each sample a dilution series was spread in 15 µl spots on NYGA plates with appropriate antibiotics and incubated for two days at 28°C. Colony-forming units (cfu) per leaf surface area were calculated for each sample. Statistical analysis of bacterial growth data was described previously (Tsuda et al., 2009). Log_10_-transformed data from all independent experiments were analyzed using the lme4 package in the R environment and the following model was fitted to the data: log_10_ cfu_gyr_ = GY_gy_ + R_r_ + e_gyr_ (GY, genotype: treatment interaction; R, biological replicate; e, residual). The mean estimates were used as the modeled log_10_-transformed bacterial titers and were compared using two-tailed t-tests. To correct for multiple hypothesis testing, the Benjamini–Hochberg method was applied.

### Quantitative real-time PCR analysis

For quantitative real-time polymerase chain reaction (qRT-PCR), plant material was harvested from leaves of the same age and developmental status as the ones used for pathogen infiltrations. Each sample included leaves from at least two plants. Total RNA was extracted from frozen plant leaves using the Gene Matrix Universal RNA Purification Kit (EURx, Gdansk, Poland). cDNA was synthesized using the High Capacity Reverse Transcription Kit (Applied Biosystems, Carlsbad, CA, U.S.A) and Oligo dT16V primer according to the manufacturer’s instructions using 1 µg RNA for a 20 µl reaction. qRT-PCRs were performed in technical triplicates on the CFX96 real-time system (Bio-Rad, Hercules, CA, U.S.A) as described previously (van Buer et al., 2016) using SYBR Green (Sigma-Aldrich, Germany) and OptiTaq Polymerase (EURx, Gdansk, Poland) and the cycling program: 95°C for 5 min, followed by 40 cycles at 95°C for 15 s, 60°C for 30 s, and finally 72°C for 30 s. All qRT-PCR primers are listed in Supplemental Table S1. The Ct values were determined using the CFX Manager software and relative expression values (ΔCt) were analyzed against the geometric mean of the *ACT2* (*Actin 2*) and the *YLS8* (*Yellow Leaf Specific protein 8*) and *RHIP* (*RGS1-HXK1 Interacting Protein 1*) transcript levels. The mean relative expression was calculated from results of three independently performed experiments. For statistical analysis, data were log_2_ transformed and analyzed using the lme4 package in the R environment (Tsuda et al., 2009): Ct_gyr_=GY_gy_+R_r_+e_gytr_ (GY, genotype:treatment interaction; R, biological replicate; e, residual). For the two-tailed t-tests, standard errors were calculated using variance and covariance values obtained from the linear model fitting. The Benjamini-Hochberg method was used to correct for multiple hypothesis testing in pairwise comparisons of the mean estimates (Tsuda et al., 2009).

### Accession numbers

Sequence data from genes described in this article can be found in the Arabidopsis Genome Initiative or GenBank/EMBL databases under the following accession numbers: *EDS1* (AT3G48090), *COR15A* (At2g42540), *ZAT10* (AT1G27730), *BAP1* (At3g61190), *PAL1* (AT2G37040), *ICS1* (AT1G74710), *PR1* (At2g14610), *NHL10* (AT2G35980), *FRK1* (AT2G19190), *tAPX* (At1g77490), *sAPX* (At4g08390), *YLS8* (AT5G08290), *ACT2* (At3g18780), *RHIP* (AT4G26410).

## SUPLEMENTAL DATA

**Supplemental Figure S1.** Bacterial titers of *Pst* DC3000 in cold-primed Col-0 and *eds1-2* 3 hours after infiltration.

**Supplemental Table S1.** List of primers used in this study.

## ACKNOWLEDGEMENTS

We thank Tina Romeis (FU Berlin/ IPB Halle) for providing *P. syringae* strains, Jane Parker (MPIPZ Cologne) for *eds1-2* mutant seeds, Ulrike Temp and Elena Reifschneider for their technical assistance.

## Notes

**Funding:** The work was supported by the German Research Foundation (CRC973/C4) and the FU Berlin.

